# Theoretical analysis of neural crest cell migration

**DOI:** 10.1101/2020.03.04.976209

**Authors:** Hamid Khataee, Andras Czirok, Zoltan Neufeld

## Abstract

The neural crest cells are embryonic stem cells that migrate throughout embryos and, at different target locations, give rise to the formation of a variety of tissues and organs. The directional migration of the neural crest cells is experimentally described using a process referred to as contact inhibition of locomotion (CIL), by which cells redirect their movement upon the cell–cell contacts. However, it is unclear how the migration alignment is affected by the motility properties of the cells. Here, we computationally model the migration alignment and the average time to reach a target location as functions of the motility dynamics and interaction of the cells in an open domain with a channel geometry. The results indicate that by varying the properties of the CIL interaction a transition takes place from random movement of the cells to an organized collective migration, where the migration alignment is maximised and the migration time is minimised. This phase transition is accelerated and strengthened with the influx rate of the cells into the domain through increasing the density of the migrating cells. The model further suggests that the migration is more coordinated when the cells with a large CIL radius move fast in a narrow domain.

## 1 Introduction

The neural crest cells are motile embryonic cells which are present in all vertebrates and produce a variety of derivatives including neurons, pigment cells of the skin, cartilage, bone, muscle, and connective tissues of the skull, face, neck, and heart [1–4]. Following the formation of neural crest cells during vertebrate embryogenesis, the cells undergo an epithelial-to-mesenchymal transition which enables them to migrate long distances along specific pathways to different target locations throughout the embryo, where they differentiate into various essential cell types [1, 4, 5]. Due to this widespread contribution of neural crest cells to nearly every major organ, these cells serve as an important model system to study the physiological and pathological processes, e.g., birth defects and invasive cancers; reviewed in [6].

The migration of neural crest cell populations is a prime representation of the collective migration of a loosely associated stream of cells, because their migration as a collective emerges from their occasional temporary interactions [7–12]. Both *in vivo* and *in vitro* experiments describe the directional migration of the neural crest cells using contact inhibition of locomotion (CIL) by which cells reshape and change their direction of movement upon the cell–cell contacts [13, 14]. CIL is also found to enhance the collective chemotaxis of neural crest cells towards a chemoattractant, where the cell density promotes the cells chemotactic response. In contrast, dispersed single cells are unable to move towards the chemoattractant [11, 15]. This presents the crucial role that CIL plays in the migration of neural crest cells during health and disease [7, 9]. For instance, the loss of CIL behaviour contributes to the malignant invasion, where the malignant tumour cells spread to the healthy tissues. This suggests that targeting the collective behaviour of a cell population may be effective in controlling the associated diseases, e.g., cancer metastasis [16]. Yet, it remains poorly understood how the dynamics of the cell-cell interactions regulate the collective migration of cells [17].

A major class of theoretical models for collective migration has been developed in the area of animal migration. These models capture the group-level migratory properties using simple interaction rules, e.g., local alignment, attraction, and repulsion [18–23]. In these models, animals are represented as point particles which interact via non-local rules, because they use sight to interact with other animals at a distance. In the context of the cell migration, however, the directionality of the migration relies on the localised information provided by direct physical contacts or chemical signals [24].

To describe the collective behaviour of neural crest cells, earlier theoretical models have used various computational techniques (e.g., agent-based and Potts models) and analytical methods (e.g., partial differential equations) [25–29]. We have also developed a two-dimensional computational model for the migratory dynamics of neural crest cells driven by CIL, in the absence of a chemoattractant [30]. Experiments have shown that the neural crest cells exhibit directional collective migration in the absence of any chemoattractant [11, 31]. We modelled cell shapes as closed contours and analysed the alignment patterns of the migration in a closed domain as functions of the cell density and shape [30].

Here, we extend our earlier work [30] by analysing the role of the CIL in the collective migration of cells in an open domain with a channel geometry, which is more realistic for the migration of neural crest cells. We also consider and compare with a simplified model based on interacting point particles describing cell-cell contacts within a certain interaction radius, without considering the cell shapes explicitly, as implemented in [18]. The results are discussed in the context of experiments and other theoretical studies.

## 2 Model description

First, we run preliminary simulations using a model based on cell contours, similar to the one described in [30]. Then, based on these observations we studied the same problem in more details using the simplified, but computationally more efficient particle based model [18].

Briefly, in [30], the migration of cells was modelled in a closed square shaped domain with periodic boundaries where the shape of the cells was initialised as contours in polar coordinates with a Gaussian function of the angle *θ*:

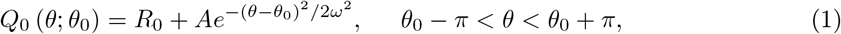

where *R*_0_ is the radius of the cell core, *A* and *ω* are the cell shape parameters, *θ*_0_ is the instantaneous cell orientation [30]. Numerically the contours are approximated by a discrete set of points. The cell shape function *Q*_0_ (*θ; θ*_0_) is then used to determine the cell velocity in the direction of net force produced by cell protrusions:

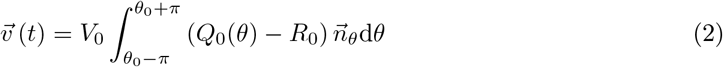

where *V*_0_ is a constant velocity parameter and 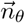 is the radial unit vector in the direction *θ*. In the absence of cell-cell interactions, a single cell moves with the velocity given by 2 with an additional random noise:

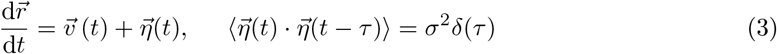

where 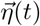 is an uncorrelated white noise with the intensity *σ*. When two cells collide, their protruded contours collapse (i.e., reset to cell core). Cells then reorient their movement by modifying the shape of their contours and move away from each other and gradually regain their normal shape [30]. The cell contours are relaxed towards the target shape function *Q*_0_ (*θ; θ*_0_(*t*)) corresponding to an isolated cell, according to:

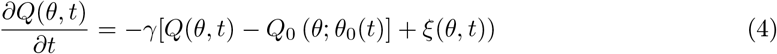

where *γ* is the regrowth rate of the cell contour; and for a more realistic representation of the cells here we also include a noise term *ξ* which describes random fluctuations of the cell protrusions, similarly to the model in [32].

Results from preliminary simulations with this cell contour model are shown in Fig. 1. Starting with an empty channel with solid reflecting boundaries at the top and bottom, the cells enter the channel on the left side at a constant rate with random initial direction. When the cells reach the right end of the channel they are eliminated from the simulation. The simulations were run until the cell numbers in the channel reached an approximate steady state. The simulation was repeated with different values of the input cell flux into the channel. The main observation from these simulations is that when the input flux is relatively high the migration is well ordered, i.e. cells move in approximately the same direction, and when the input flux is low there is a lower cell density in the channel and the cells often loose the overall direction of the migration and follow a longer random forward-and-backward trajectory to get through the channel; see Fig. 1(a, b) and Movies 1 and 2. The collective alignment of migrating cells can be characterised by the order parameter:

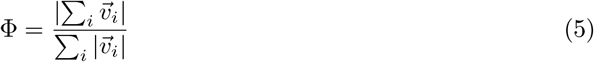

where Φ varies between 0 (uncorrelated random movement) and 1 (fully aligned migration). The order parameter as a function of time for two values of the input flux is shown in Fig. 1(c).

**Figure 1:**
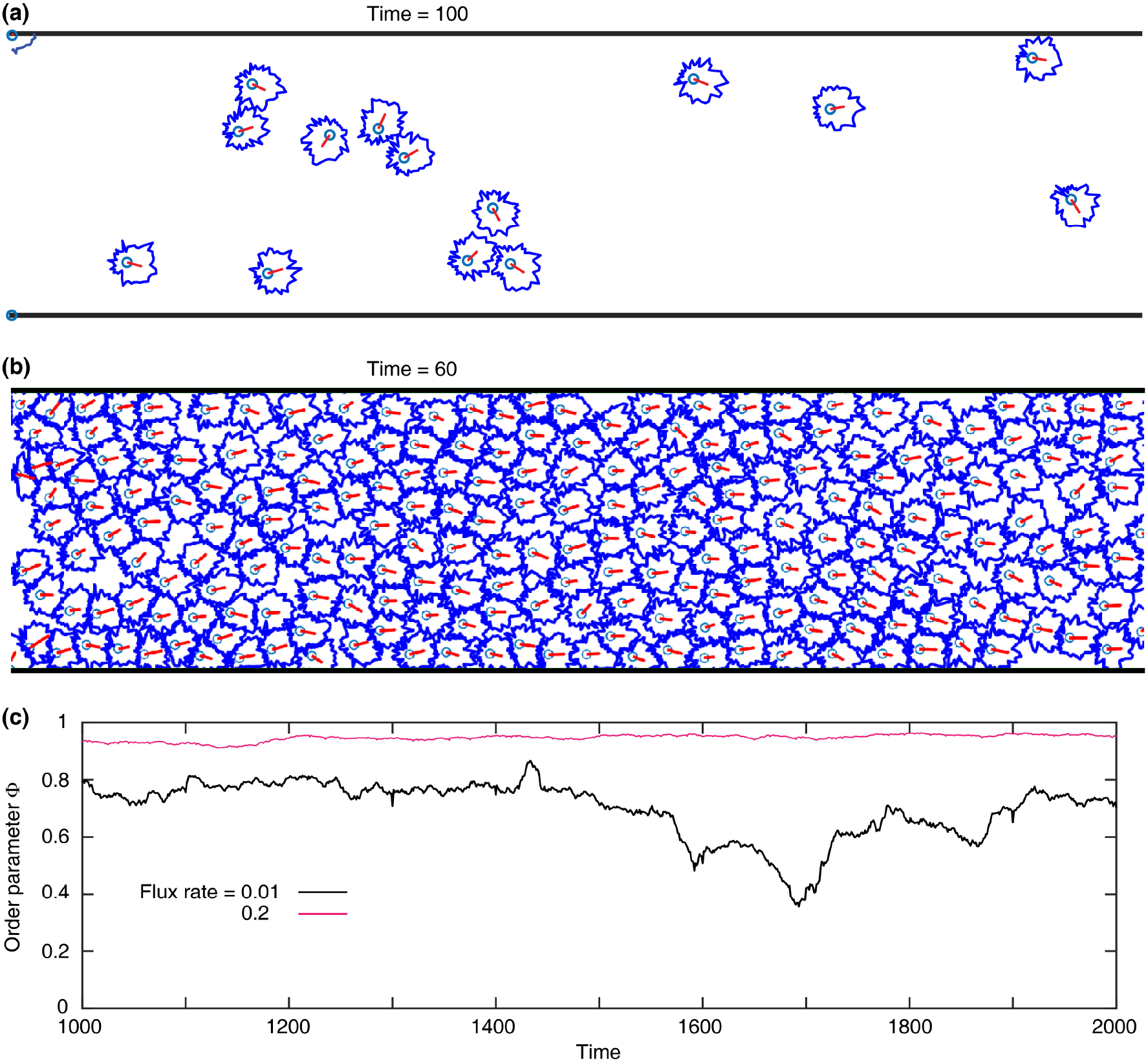
Migration of cells with contour shapes. (a, b) Snapshots of cells which in an open migratory domain with a channel geometry and dimensions *L_x_* = 40 and *L_y_* = 10 in length-units, where top and bottom boundaries are reflective. Flux rates are 0.01 (a) and 0.2 (b). Other simulation parameters are: *R*_0_ = 0.5, *A* = 0.5, *ω* = 1, *V*_0_ = 0.3, and *γ* = 0.1. For each cell shape, red arrow: the instantaneous cell velocity; light blue circle: cell body with radius *R*_0_; dark blue contour: cell shape; see Equation (1). (c) Order parameter Φ versus time for the simulations in (a) and (b).

In order to characterise the properties and more detailed parameter dependence of the cell migration in the open channel, we use a simplified particle-based computational model. We have shown earlier [30] that in a closed domain the contour-based cell model has similar properties and produces a qualitatively similar phases of ordered coherent and irregular movement as the self-propelled particle model introduced in [18]. Therefore, we model the migratory properties of particle-based cells in an open domain.

We modify the closed domain of the original model by assuming that the self-propelled particles enter the domain on the left side of the channel at a fixed rate *f*, defined as the number of new cells added per unit time (see Movie 3). The model is discrete in time so the time step taken to update the cell positions is Δ*t* = 1. Following [18], each cell *i* moves in the domain by updating its position 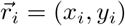 at each time step as:

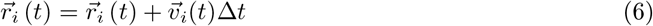

with velocity of magnitude *v*_0_:

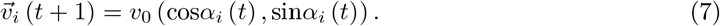

The direction of the displacement is calculated as

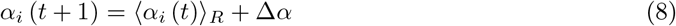

where 〈*α_i_*(*t*)〉_*R*_ is the average of the velocity directions *α_i_* of the cells (including cell *i*) within an interaction circle of radius *R* surrounding cell *i*. Δ*α* is a random noise term uniformly distributed over the interval [–*η*/2, *η*/2] [18]. Finally, when a cell reaches the right end of the channel it is removed from the simulation, resembling a target region where neural crest cells stop migrating and differentiate into various cell types [25, 33].

Since the cells are moving in an open domain, the cell density *ρ* is variable, not prescribed as in the closed system studied earlier [18, 30], and is determined by the influx rate of the cells. By simplifying the cell shapes into a single cell-cell interaction radius, the number of free parameters is reduced, having only the flux parameter *f* in addition to the three cell parameters *v*_0_, *η*, and *R*. Using the simulations based on Equations (6–8), we focus on three characteristics of the collective migration: the alignment of migration calculated using the order parameter defined in Equation (5), the migration time, and the cell density in the domain. The migration time *t_m_* for a cell is defined as the time taken for crossing the channel from left to right. Finally, the cell density *ρ* is calculated by binning the domain along the *x*-dimension, where the width of each bin is five length-units. The cell density *ρ* is then calculated as the average number of cells located in each bin divided by the width of the domain *L_y_*.

## 3 Results and discussion

We start with exploring how the migratory characteristics Φ, *ρ*, and *t_m_* change over time. We find that increasing the flux rate *f* enhances the migration alignment, accelerates the migration time, and increases the cell density in the domain; see Figure 2(a-d).

**Figure 2:**
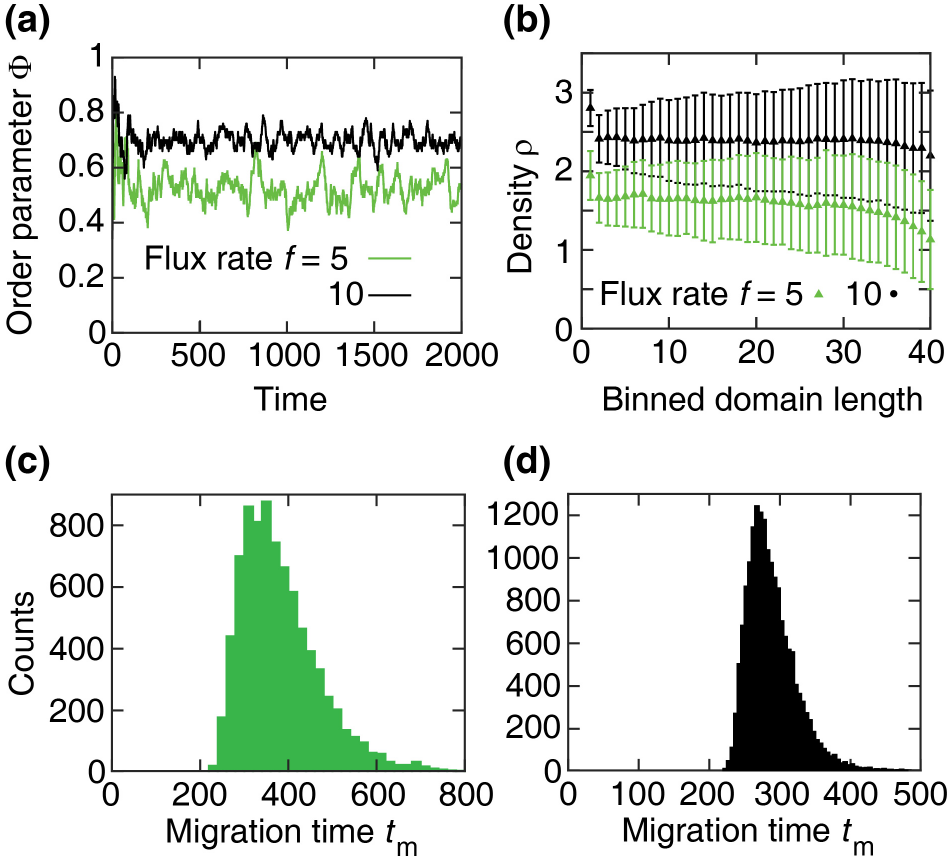
Migratory dynamics over time. (a) Order parameter Φ versus time. For *f* = 5, Φ = 0.53 ± 0.05. For *f* = 10, Φ = 0.70 ± 0.04. (b) Cell density *ρ* versus the binned length of the domain, where each bin is five-length-unit wide. The steady *ρ* are 1.58 ± 0.14 and 2.40 ± 0.08 for flux rates *f* = 5 and 10, respectively. (c, d) Migration time *t*_m_ versus the number of cells exited the domain. (c) For *f* = 5, *t*_m_ = 381.32 ± 89.02. (d) For *f* = 10, *t*_m_ = 287.34 ± 33.69. Other simulation parameters are *v*_0_ = 1, *η* = 1, and *R* = 0.5.

We then examine these migratory characteristics in response to changes in the cell’s parameters. In addition, we analyse the effect of the width of the domain *L_y_* on these characteristics, since earlier studies have shown that a spatial confinement [27] and growing domain [24, 34] may affect the collective migration of cells. Results in Fig. 3(a-c) show that increasing the interaction radius enhances the migration alignment and accelerates the migration time, while having little effect on the cell density. Interaction radius R leads to a transition from random movement of cells (low Φ) to an organized collective flow (high Φ), i.e., there is a rise in the alignment of the cells movement (and drop in the migration time) as R approaches a critical value above which the migration alignment (and time) remains steady. This agrees with the calculations of our earlier model [30] that cells with broader contours produce better migration alignment. The model further predicts that the critical interaction radius depends on the flux rate, suggesting that CIL can dramatically enhance the migration alignment when the cells enter a migratory pathway with a high influx rate. At lower influx rates, a coordinated and fast collective migration would require a larger interaction radius.

**Figure 3:**
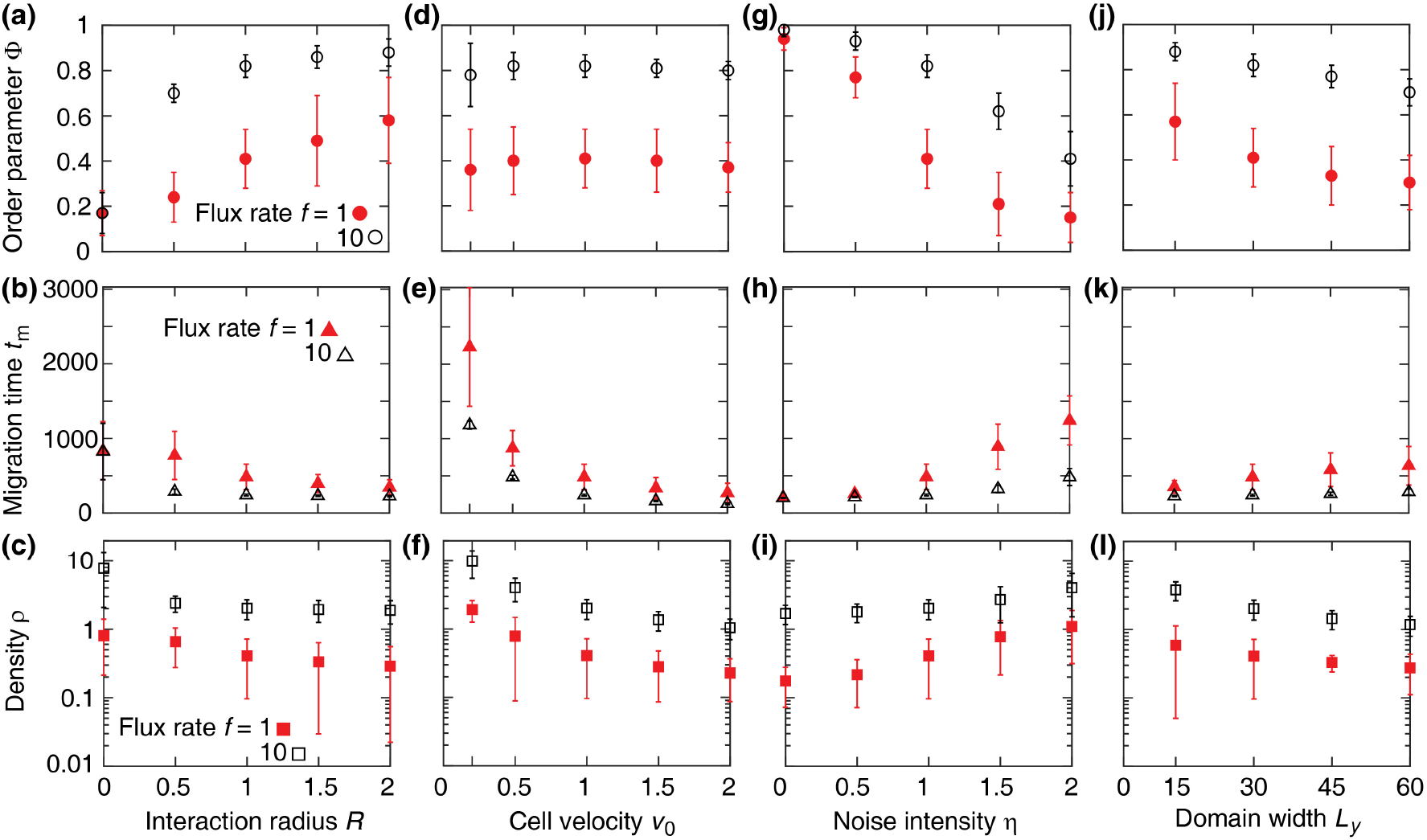
Migratory dynamics. Order parameter Φ (top row), migration time *t*_m_ (middle row), and cell density *ρ* (bottom row) versus the interaction radius *R* (a-c), cell velocity *v*_0_ (d-f), noise intensity of the cell motility *η* (g-i), and width of the domain *L_y_* (j-l). In (a-i), *L_y_* = 30. In (a-c), *v*_0_ = 1 and *η* = 1. In (d-f), *η* = 1 and *R* = 1. In (g-i), *v*_0_ = 1 and *R* = 1. In (j-l), *v*_0_ = 1, *η* = 1, and *R* = 1.

The cell velocity parameter *v*_0_ is found to slightly increase the migration alignment, and lowers the migration time and cell density as shown in Fig. 3(d-f). When cells move faster, the frequency of the cell-cell collisions increases, suggesting that the cell velocity enhances the CIL effect on coordinating the cell migration. Together, results in Fig. 3(a-f) suggest that faster cells with greater interaction radius would enhance the the migration alignment and accelerate the migration time.

The analysis of the stochasticity of the cells movement indicates that the noise intensity *eta* can transition a coordinated migration to a random movement of the cells. The randomness in the movement of the cells blocks their coordination in migrating as a collective. In turn, this would increase the cell density in the domain; see Fig. 3(g-i).

Finally, we find that the collective migration becomes more aligned and faster with narrowing the domain; see 3(j-l). A narrow domain increases the chance of cell collisions, suggesting that the spacial confinements would strengthen the effect of CIL in coordinating the collective migration.

We now examine the migratory dynamics Φ, *t_m_*, and *ρ* in response to the influx rate *f* of the cells into the domain. We find that the flux rate transitions a random movement to an organized migration, where the migration alignment is maximised and the migration time is minimised; see Fig. 4(a, b). A similar transition from a sparse domain into a dense is calculated with increasing *f*; see Fig. 4(c). This later result allows us to examine the phase transitions in *φ* and *t_m_*, in Fig. 4(a, b), as a function of *ρ*. This way, we can compare our results directly with the calculations of the cell-contour model from [30].

**Figure 4:**
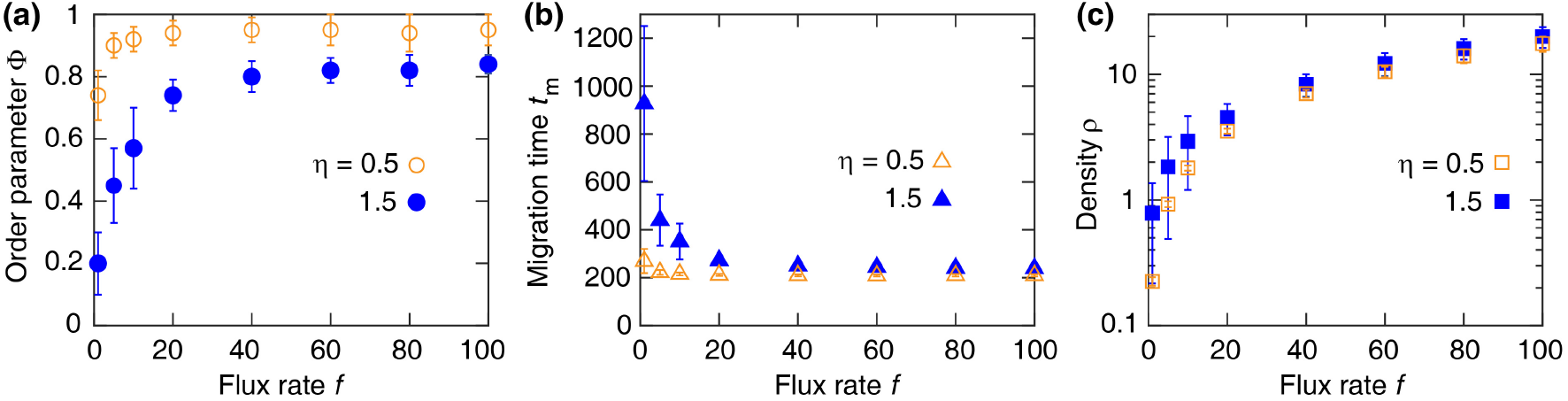
Migratory dynamics versus the influx rate *f*. Order parameter Φ (a), migration time *t*_m_ (b), and cell density *ρ* (c) calculated with model parameters *v*_0_ = 1and *R* = 0.9 for *η* = 0.5 and 1.5.

The density-dependent analysis of the migration dynamics indicates that the cell density transitions the uncoordinated movement of cells to a coordinated migration, where the migration time is minimised; see Fig. 5(a, b). To compare these results with our earlier results that considered the cell shapes as contours [30], we repeat the simulations in a closed domain, as performed in [30]. Regardless of the domain boundaries (open or closed), we find the same phase transition in Φ in response to the cell density *ρ*; see Fig. 5(a, c). This phase transition is also consistent with the phase transition obtained by considering the cell contour shapes, showing that our simplified model can reproduce the results obtained by the more complex model in [30]; compare Fig. 5(c, d).

**Figure 5:**
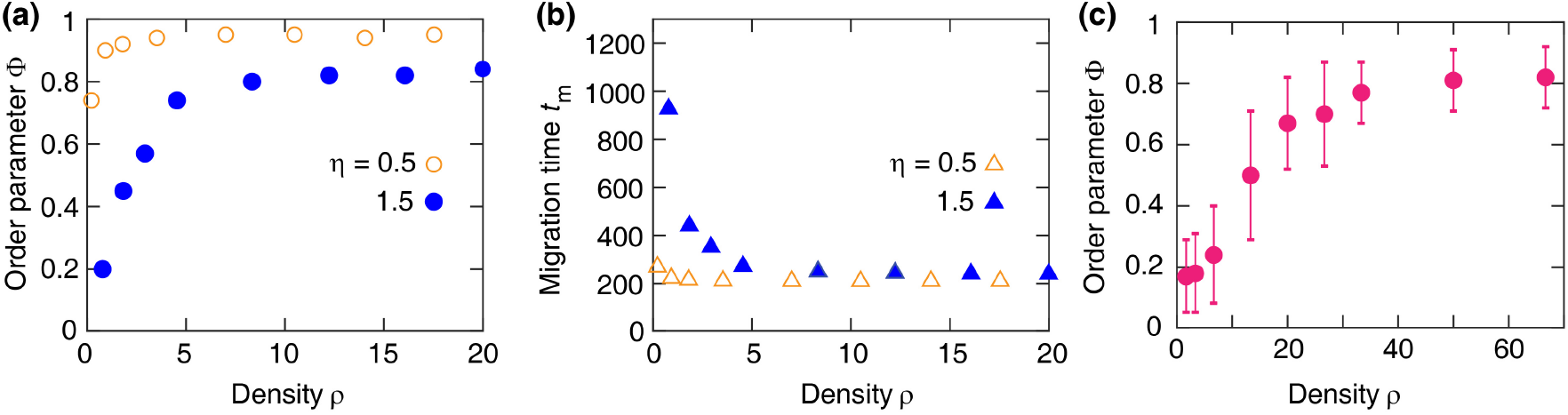
Comparison of the models. (a, b) Order parameter Φ and migration time *t*_m_ versus the cell density *ρ* calculated with parameters *v*_0_ = 1 and *R* = 0.9 for *η* = 0.5 and 1.5. (c) Φ when the domain is square (30 × 30, in length units) with periodic boundaries in x-dimension, as performed in [18]. Simulation parameters are as in (a).

The results of our computational model show several agreements with the experimental and earlier theoretical findings. Experiments have shown that the CIL dramatically increases the percentage of the neural crest cells that reach their target region over time [25, 27]. Likewise, Woods et al. [26] found that the cell-cell interactions reduce the time taken for the cells to reach a target location. These observations confirm our calculations for the migration alignment and time in Fig. 2(a, b). The analysis of the effect of the randomness in the movement of the neural crest cells in Fig. 3(g-i) has shown that stochastic fluctuations in the movement of cells limit the coordination of the migration. This accords with our earlier calculations in [30], the flocking models (where the self-propelled particles had non-local interactions) [18, 35], the self-propelled short-range interacting agents in swarm migration [36], and the self-propelled interacting cells [37] that the amplitude of the stochasticity of the movement of cells leads to random movement.

In addition, previous works on the effect of the spacial confinements on the migration of neural crest cells have shown that removing the lateral confinement of the domain significantly reduces the directionality of the collective movement [27]. In agreement with these findings, the model calculated higher order parameter Φ for narrower domains; see Fig. 3(j-l).

Finally, *in vivo* experiments have shown that, when the neural crest cells contact during migration, their migratory persistence (i.e., the maximum distance along a cell trajectory divided by the total length of that trajectory) increases [27]. Moreover, the self-propelled particle models have indicated that the particle density transitions particles’ random movement to a coordinated migration [35, 36]. Together, this implies that the coordination of the migration would enhance in response to an increase in the cell density in the domain. This has been shown in our density-dependent calculations in Fig. 5(a-c).

In conclusion, we have an extended computational model to analyze the migration of the neural crest cells. The model has extended our earlier analysis in a closed domain [30] to an open domain. This allowed us to take into account the effect of the flux rate of the cells into the domain on the migration alignment and time. It also helped to calculate the cell density in the domain. These benefits are gained along with faster computational simulations, as the cell-cell interactions are modelled using an interaction radius.

## Supporting Information

### Movie 1.

Collective neural crest cell migration an open domain with dimensions *L_x_* = 40 and *L_y_* = 10, where cell shapes are modelled as closed contours. Flux rate is 0.01. Other simulation parameters are: *R*_0_ = 0.5, *A* = 0.5, *ω* = 1, *V*_0_ = 0.3, and *γ* = 0.1. For each cell shape, red arrow: the instantaneous cell velocity; light blue circle: cell body with radius *R*_0_; dark blue contour: cell shape; see Equation (1).

### Movie 2

Collective neural crest cell migration an open domain with dimensions *L_x_* = 40 and *L_y_* = 10, where cell shapes are modelled as closed contours. Flux rate is 0.2. Other simulation parameters are: *R*_0_ = 0.5, A = 0.5, *ω* = 1, *V*_0_ = 0.3, and *γ* = 0.1; see Equation (1).

### Movie 3

Collective neural crest cell migration an open domain with dimensions *L_x_* = 200 and *L_y_* = 30, where cells are modelled as self-propelled particles. Simulation parameters are: *v*_0_ = 1, *η* = 1.5, *R* = 0.9, and *f* = 10.

